# Performance of Complex Visual Tasks using Simulated Prosthetic Vision via Augmented-Reality Glasses

**DOI:** 10.1101/707851

**Authors:** Elton Ho, Jack Boffa, Daniel Palanker

## Abstract

**Purpose:** Photovoltaic subretinal prosthesis is designed for restoration of central vision in patients with age-related macular degeneration (AMD). We investigated the utility of prosthetic central vision for complex visual tasks using augmented-reality (AR) glasses simulating reduced acuity, contrast and visual field.

**Methods:** AR glasses with blocked central 20° of visual field included an integrated video camera and software which adjusts the image quality according to three user-defined parameters: resolution, corresponding to the equivalent pixel size of an implant, field of view, corresponding to the implant size, and number of contrast levels. The real-time processed video was streamed on a screen in front of the right eye. Nineteen healthy participants were recruited to complete visual tasks including vision charts, sentence reading, and face recognition.

**Results:** With vision charts, letter acuity exceeded the pixel-sampling limit by 0.2 logMAR. Reading speed decreased with increasing pixel size and with reduced field of view (7-12°). In the face recognition task (4-way forced choice, 5° angular size) participants identified faces at >75% accuracy, even with 100 μm pixels and only 2 grey levels. With 60 μm pixels and 8 grey levels, the accuracy exceeded 97%.

**Conclusions:** Subjects with simulated prosthetic vision performed slightly better than the sampling limit on the letter acuity tasks, and were highly accurate at recognizing faces, even with 100 μm/pixel resolution. These results indicate feasibility of the reading and face recognition using prosthetic central vision even with 100 μm pixels, and performance improves further with smaller pixels.

## Introduction

Age-related macular degeneration (AMD) is a leading cause of untreatable visual impairment. With the current prevalence of 8.7% worldwide, AMD is projected to affect almost 200 million people in 2020, and its prevalence is growing with the population aging (1, 2). Patients with advanced atrophic AMD (currently about 1% prevalence in Western countries (1, 2)) suffer from the loss of photoreceptors in the macula, leading to compromised central vision. Although high-resolution vision is lost, patients still can use their preserved peripheral vision and typically retain acuity no worse than 20/400. Therefore, restoration of central vision may be worthwhile only if the restored visual acuity exceeds the residual natural level.

In the healthy eye, photoreceptors convert incident light into electrical and chemical signals. The resultant neural signals are processed by the bipolar cells and other non-spiking neurons in the inner nuclear layer (INL) and advance to the retinal ganglion cells (RGC), which generate action potentials that propagate via optic nerve to the brain. Loss of photoreceptors in retinal degenerative diseases impairs the initial phototransduction process, while the remaining retinal network remains intact, albeit with some rewiring (3–5).

Multiple approaches are being developed to address the loss of sight in the retinal degeneration(6), including gene therapy (7), cell transplantation (8, 9), optogenetics (10), and electronic implants. In the latter case, an array of electrodes is placed at the stimulation site, such as the retina (11), optic nerve (12), lateral geniculate nucleus (LGN) (13), or primary visual cortex (14). Electric current is injected into tissue to stimulate cells and thereby elicit visual perception. Upon electrode activation, patients report perceiving “bright spots”, termed phosphenes (15, 16). The number of electrodes limits the amount of information deliverable, and electrode density restricts the highest possible resolution. In animal studies with photovoltaic retinal prosthesis, we demonstrated that grating acuity matches the pixel pitch with 55 (17) and 75μm pixels (18). Recent clinical trial of such implants (PRIMA, by Pixium Vision) having 100μm pixels also demonstrated that prosthetic visual acuity in AMD patients is only 10-30% below the sampling limit of 20/420 for the current pixel size (19).

This subretinal implant stimulates the first layer of neurons after photoreceptors (INL), and therefore elicited network-mediated retinal responses retain many features of the natural signal processing, including flicker fusion at high frequencies (>20 Hz) (18, 20), adaptation to static images (21), antagonistic center-surround organization of receptive fields with linear and nonlinear summation of its subunits (22). Patients with the PRIMA implant can perceive lines of a single pixel in width, and identify letters with the minimum gap in the letter C of 1.1 – 1.3 pixels (19).

Since AMD patients retain peripheral vision, they have little problem with ambulation. However, impaired reading and face recognition pose significant challenges in daily living (23). To assess the spatial resolution, number of levels of grey and the size of the implant required for these visual tasks, we simulated prosthetic central vision using augmented-reality glasses with a camera. Prosthetic vision was mimicked by controlled reduction in spatial resolution, contrast and visual field of the images projected on the built-in display. Current clinical version of the PRIMA subretinal array has pixel pitch of 100 μm (19). In rodents, we already validated feasibility of 75 (18) and 55 μm pixels (17), and they may be further reduced down to 20 μm using 3-dimensional electrodes (24). Here, we investigate how well healthy subjects can accomplish complex visual tasks, including reading and face recognition, under various levels of image degradation.

Studies of simulated prosthetic vision were conducted in the past, but we find those results insufficient for predicting the outcomes with our current implant. With photovoltaic subretinal implant for restoration of central vision in AMD patients, simulation requires the following specifications: (a) pixel density >100 pixels/mm^2^, (b) no gaps between phosphenes, (c) visual field in the range of 7-10°, and (d) eye scanning is allowed. Since previous studies did not address these specifications, we conducted a psychophysics study to assess the limits of visual performance of the PRIMA system and set the expectations for the upcoming clinical trials.

## Methods

### Subjects

Nineteen subjects, all recruited from personnel at Stanford University, signed informed consent and participated in the current study. All subjects had self-reported normal vision, and their visual acuity was verified with both a Landolt C test and ETDRS chart prior to the experiments. For complex reading tasks, subjects were required to have native or near-native English proficiency. All subjects had limited or no prior experience with virtual- or augmented-reality (AR) glasses. The study was approved by the Stanford IRB panel on human subjects research and conducted according to the institutional guidelines, following the tenets of the Declaration of Helsinki.

### Experimental setup

The experimental apparatus included two parts: a stimulus presentation system and AR glasses with the head-on display and an image processing unit (Figure 1).

**Figure 1.**
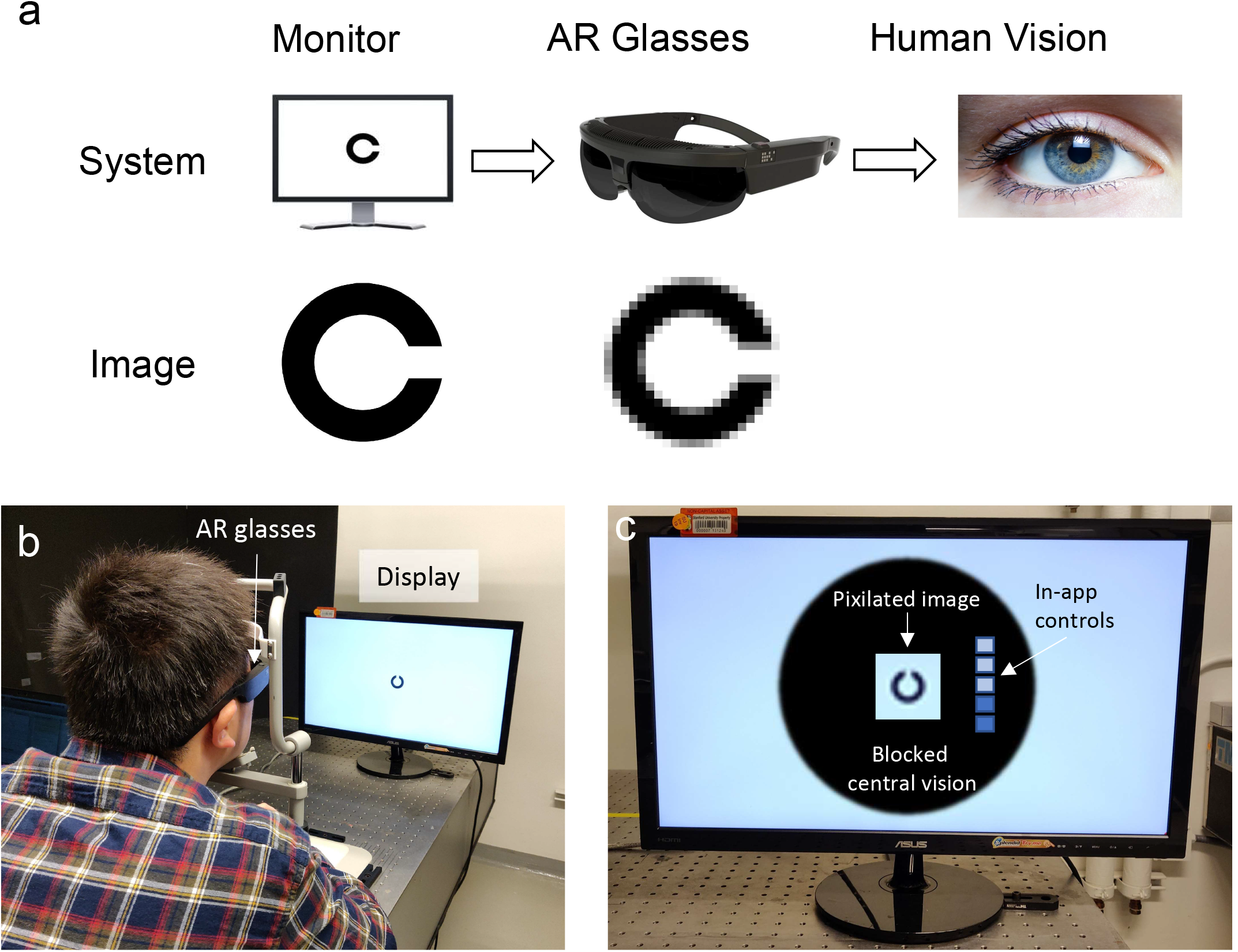
Experimental setup. (a) Schematic of the experimental setup. High resolution images are presented on a monitor. The front camera of the augmented-reality (AR) glasses captures the video stream. Custom software pre-loaded on the AR glasses adjusts the video quality to mimic prosthetic vision and displays it in the AR glasses. (b) A subject in front of the apparatus. (c) Illustration of vision through the AR glasses.

The stimulus presentation system involved a 24” monitor (ASUS VS248H-P) controlled by a laptop computer (Thinkpad 25, Lenovo) using a PsychToolbox-based (25–27) custom software in Matlab. This system was used to display stimulus and record subjects’ responses (such as accuracy and time taken) via experimenter input. The monitor was placed 30” away from a chinrest, where the subjects would place their head during an experiment. The monitor had resolution of 2400 x 1350 pixels, corresponding to 90.4 pixels per degree of visual angle (ppd).

Camera (4 MP) mounted on the front of the AR glasses (ODG R-7, Osterhout Design Group, San Francisco, CA) captures a live video stream. Camera magnification was set to match the angular size of the natural vision. The data is then processed with an Android-based custom app in real time according to three user-defined parameters: pixilation (equivalent to 30-100 μm pixel size on the retina), number of grey levels (2–256), and field of view (FOV) (7-12 degrees). The resultant video was presented on the display in the glasses (specs: 30° FOV, 720p, 80 fps). The latency between the camera and the display was minimal due to fast video processing. In a typical AR display system, the integrated display is transparent, so that presented visual information can fuse with the passthrough background (hence, “augmented” reality). To mimic vision loss in AMD patients, an area on the glasses corresponding 20° of central vision, was blocked with black opaque tape for both eyes. In this region, only the integrated display was visible, while outside that region, only natural peripheral vision was present (Figure 1c). Here we only assess monocular prosthetic vision, so the display was only switched on for the right eye, which incidentally corresponded to the dominant eye of all subjects.

The video processing was done with the OpenCV library, and the workflow was as following: A video frame was cropped to match the desired FOV. The frame was then converted to greyscale, downsized, and re-upsized back to the original image size, resulting in a tightly-packed pixilated greyscale image. We used the default nearest-neighbor interpolation for image transformation in Android. The pixilation here matched the desired pixel size on the retina, e.g. 100 μm pixels subtend 0.35° on the human retina. The grey color value for each pixel was then rounded to the nearest 255/*n*, where *n* is the number of contrast levels.

Subjects were instructed to wear the AR glasses and learned to adjust pixel size, contrast levels, and FOV using in-app controls. To familiarize our subjects with simulated prosthetic vision, they were instructed to look around the laboratory freely for a few minutes, and were also presented pictures of common animals, plants, and foodstuffs.

### Procedures

We conducted three different experiments: (1) single-character visual acuity (SCVA), (2) sentence reading, and (3) face recognition. Parameters for simulated prosthetic vision are summarized in Table 1. Subjects were instructed to fixate their central vision to the center of the AR screen, but were allowed to move their eyes and head, if desired. In all experiments, subjects vocalized their responses, which were recorded and timed by the experimenter. Typically, a full set of experiments could last up to 90 minutes. If a subject got tired, a new session for remaining tasks was scheduled.

**Table 1.**
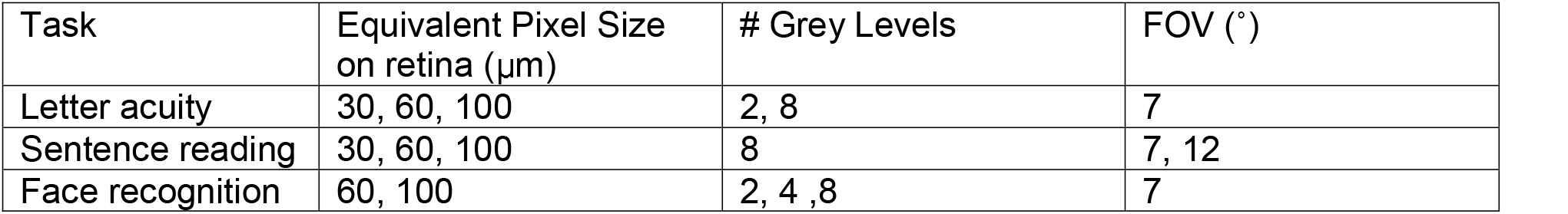
Parameters of the image processing used for each experiment.

| Task | Equivalent Pixel Size on retina ($\mu\text{m}$) | # Grey Levels | FOV ($^\circ$) |
| --- | --- | --- | --- |
| Letter acuity | 30, 60, 100 | 2, 8 | 7 |
| Sentence reading | 30, 60, 100 | 8 | 7, 12 |
| Face recognition | 60, 100 | 2, 4, 8 | 7 |

### Letter acuity

Subjects (n=13 for 30 and 60 μm pixels; n=19 for natural vision and 100 μm pixels) were asked to identify the orientation of the Landolt C, presented one at a time. If the subject could identify at least four out of five orientations of the same size, we reduced the letter size by 0.1 LogMAR units and repeated. The same experiment was conducted also with ETDRS letters in Sloan font. The smallest feature of these characters was 1/5^th^ of the letter size. Subjects were first tested for their visual acuity with normal or corrected-to-normal vision without AR glasses, and then with simulated prosthetic vision. As a point for comparison, we also computed the sampling limit for each prosthetic pixel size by calculating its geometric-equivalent visual acuity.

### Sentence reading

Subjects (n=9 for simple sentences; n=10 for complex sentences) were asked to read aloud displayed sentences as fast as possible, following standard MNREAD protocol (http://legge.psych.umn.edu/mnread-set). Text in Arial font was presented in three lines, with approximately 20 characters per line. A new sentence with reduced font size (−0.1 LogMAR) was displayed upon successful utterance (≤2 mistaken words per sentence). The font size was measured as the visual angle between the top of the letter “k” and the bottom of the letter “p”. In between the sentences, a fixation cross was shown in the center of the screen for 2 seconds to re-center the subjects’ vision. Subjects were first tested with their normal/corrected binocular vision, and then with simulated prosthetic vision with varying pixel size and FOV. The reading speed (in words per minute, or WPM) for each sentence was recorded in software. We evaluated reading performance on three key metrics: reading acuity (RA, smallest resolvable sentence), maximum reading speed (MRS), and critical font size (CFS, smallest font size at which MRS is reached).

The texts used can be classified into simple and complex sentences. Simple sentences were either composed in-house according to the MNREAD protocol, or taken from the MNREAD iPad App ©2017 (https://itunes.apple.com/us/app/mnread/id1196638274?ls=1&mt=8). Complex sentences were selected from the Manually Annotated Sub-Corpus (MASC) from the Open American national Corpus (OANC) (http://www.anc.org/) with three criteria: number of characters between 55 and 70, average word length between 5.5 and 6.5, and the sentence can be segmented into three lines of similar length. Generally, simple MNREAD sentences have stand-alone context and involve vocabulary at elementary school level in the US (e.g. “He looked up at his mother and told her he was really happy”), while complex sentences may incur more context with advanced vocabulary (e.g. “Good housekeeping contributes to safety and reliable results”). Only subjects with native or near-native English level were selected for the complex reading task. Results for simple and complex sentences were cross-compared using a two-sample t-test.

### Face recognition

Subjects (n=19 for 100 μm; n=17 for 60 μm) were shown a reference adult face and required to select one out of four other faces that matched the identity of the reference (Figure 2a). The correctness and time taken for each selection were recorded. A set of 10 trials were performed for each parameter combination, which were presented in a pseudorandom order to minimize learning effects.

**Figure 2.**
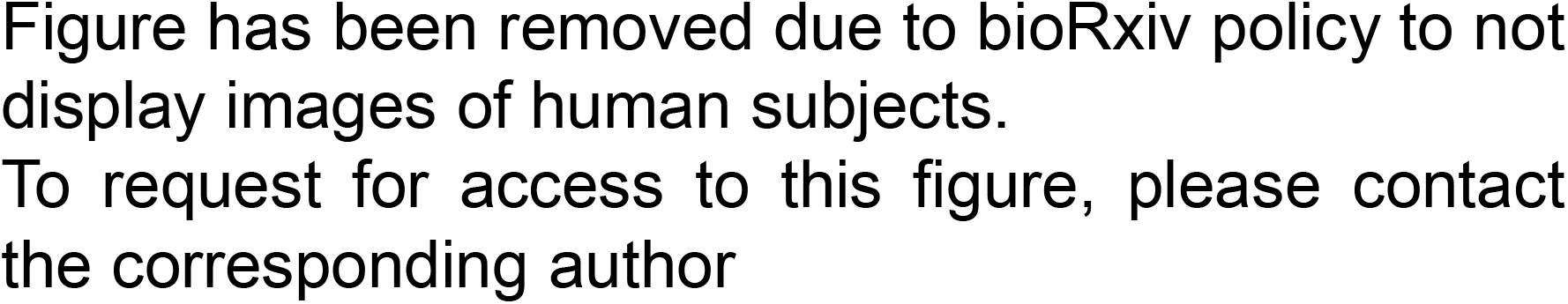
Face recognition task. (a) An example set of five faces presented. Subjects were asked to pick the face that matches the identity of the central person. Each face spanned approximately 5°x5°. (b) Effects of the number of grey levels and resolution on an image.

Images of non-occluded adult heads were randomly selected from the Face Place database (http://www.tarrlab.org/). The database is licensed under a CC BY-NC-SA 3.0 Unported License. For the same identity, a set of images included different viewing angles and facial expressions, with the background cropped out. Generally, the most prominent features above the neck were visible, including hair style, skin tone, and both eyes. Images were resized and cropped to 5°x5°. Five images were tiled as shown in Figure 2a, occupying a visual field of 16°x16°. The reference image was placed at the center.

## Results

### Letter acuity

With both ETDRS letters and Landolt C, VA improved with reduced pixel size, as shown in Figure 3. VA measured by both testing paradigms agree with each other. Landolt C test yielded slightly better VA than ETDRS letters - by 0.05 logMAR, albeit insignificant (Supp. Figure 1). Decreasing contrast level from 8 to 2 did not affect VA significantly. All measured VA were at least 0.2 logMAR higher than the computed sampling limit for each pixel size. This could be attributed to oversampling by scanning and subjects looking for differences between undersampled letters. Letter recognition here required around 3 pixels per letter width for all pixel sizes, agreeing with the 3 – 7 phosphenes per letter width reported by other studies (28–30).

**Figure 3.**
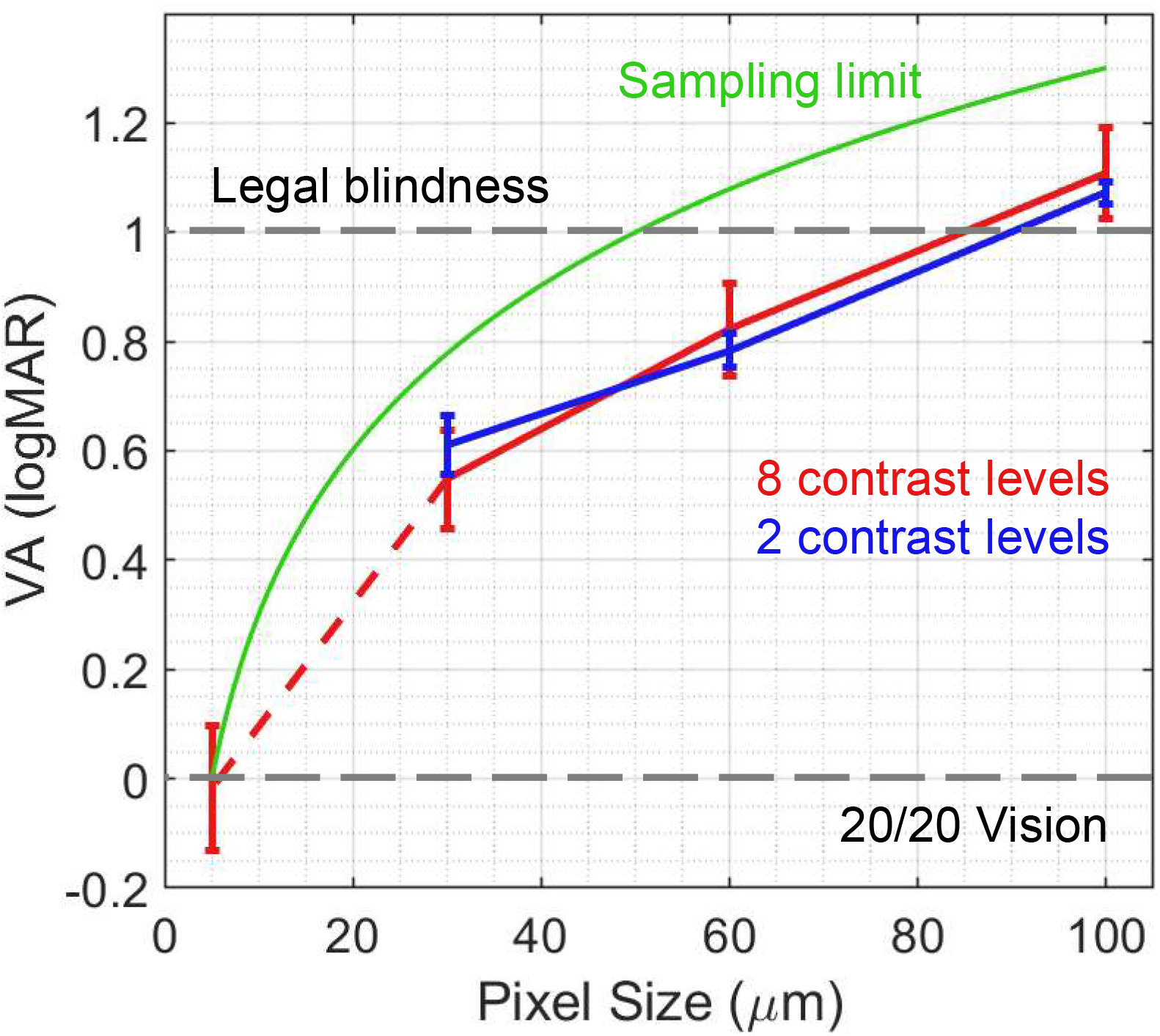
Letter acuity results (n=13 for 30 and 60 μm pixels; n=19 for natural vision and 100 μm pixels). The leftmost data point at 5 μm indicates VA for natural vision of the subjects. Error bars are presented in terms of s.d.

Most subjects self-reported that near the limit, they did not explicitly resolve the opening of a Landolt C. They employed a strategy where they scanned the object and identified the side of the blob that flickered more, through which correctly determining the orientation.

### Sentence reading

With limited pixel size and FOV, reading speed with simulated prosthetic vision (Figure 4, green and red lines) was much slower than that with unobstructed natural vision (blue line). Reading acuity (RA) for both natural and prosthetic vision matched the corresponding letter acuity. As the font size increased above VA threshold, reading speed rapidly increased until the maximum reading speed (MRS) was reached at the critical font size (CFS). Further increase of the font size was detrimental, as fewer words and letters could fit in the FOV. For example, a nine-letter word of 1.5° font size (corresponding to 1.5° vertical height and 0.78° horizontal width allotted to each letter) can barely fit into 7° FOV. For all pixel sizes, CFS was around double the RA, and the smallest readable font size was about 2.5 pixels per letter width, slightly less than the letter acuity test and previous reports. The discrepancy can be attributed to the fact that in reading tasks, the loss in letter-by-letter information is compensated by contextual clues.

**Figure 4.**
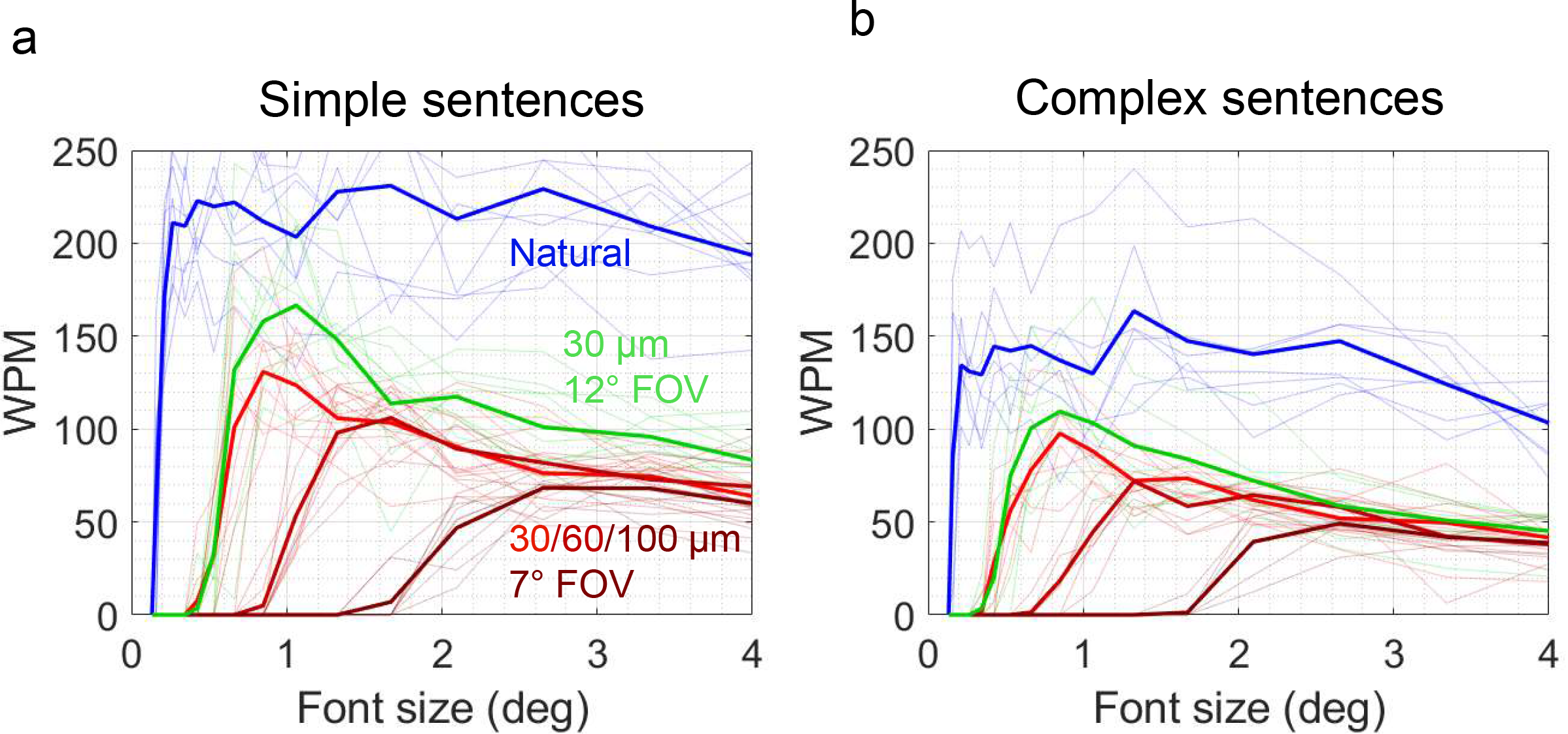
Sentence reading speed. (a) Simple sentences. (b) Complex sentences. Faded lines represent individual measurements, and the bold lines represent the population mean.

**Figure 5.**
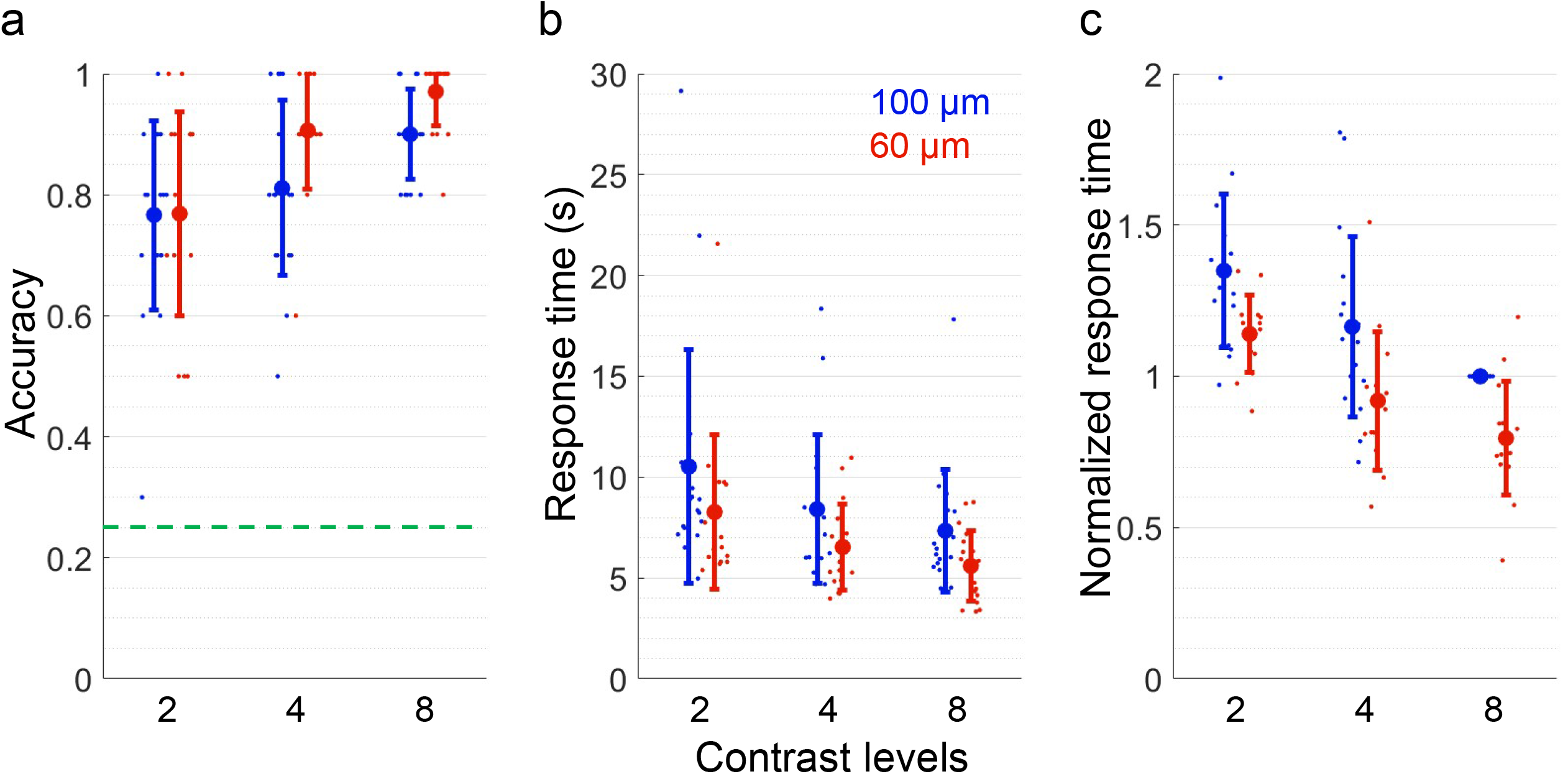
Face recognition. (a) Accuracy. (b) Response time. (c) Response time normalized to 100 μm pixels and 8 grey levels. Each dot represents an independent measurement. Error bars are presented in terms of s.d.

Generally, smaller pixels allowed for denser sampling, resulting in better RA, MRS, and CFS. Meanwhile, an increased FOV did not significantly affect RA, while raising reading speed with all font sizes greater than CFS (*t*=3.2, *p*=0.005 for MRS). The numerical results are summarized in Table 2.

**Table 2.** Reading acuity (VA), maximum reading speed (MRS), and critical font size (CFS) for reading MNREAD sentences using simulated prosthetic vision. All errors are reported as standard deviation.

| Parameters (pixels/FOV) | RA (deg) | MRS (WPM) | CFS (deg) |
| --- | --- | --- | --- |
| 100 $\mu\text{m}$ / $7^\circ$ | $1.60 \pm 0.15$ | $58 \pm 10$ | $3.0 \pm 0.6$ |
| 60 $\mu\text{m}$ / $7^\circ$ | $0.81 \pm 0.08$ | $77 \pm 16$ | $1.7 \pm 0.7$ |
| 30 $\mu\text{m}$ / $7^\circ$ | $0.46 \pm 0.10$ | $106 \pm 22$ | $1.0 \pm 0.6$ |
| 30 $\mu\text{m}$ / $12^\circ$ | $0.45 \pm 0.06$ | $141 \pm 27$ | $1.1 \pm 0.2$ |
| Natural | $0.14 \pm 0.01$ | $201 \pm 40$ | $0.36 \pm 0.07$ |

General trends with complex sentences were the same, albeit at lower speed. However, the effect of FOV on MRS became insignificant (e.g. *t*=1.45, *p*=0.156 for 30 μm/12°). Counterintuitively, RA and CFS were slightly better (smaller) for complex sentences than for simple ones, possibly due to the word predictability in context-rich sentences.

**Table 3.** Reading acuity (RA), maximum reading speed (MRS), and critical font size (CFS) for reading complex sentences using pixelated vision. All errors are reported as standard deviation. Asterisk (*) indicates *p*<0.05 (2-sample *t*-test) compared to simple sentences with the same parameters.

| Parameters (pixels/FOV) | RA (deg) | MRS (WPM) | CFS (deg) |
| --- | --- | --- | --- |
| 100 $\mu\text{m}$ / $7^\circ$ | $1.63 \pm 0.11$ | $52 \pm 7$ | $2.7 \pm 0.4$ |
| 60 $\mu\text{m}$ / $7^\circ$ | $0.71 \pm 0.11^*$ | $73 \pm 10$ | $1.7 \pm 0.5$ |
| 30 $\mu\text{m}$ / $7^\circ$ | $0.36 \pm 0.05^*$ | $102 \pm 18$ | $0.80 \pm 0.15$ |
| 30 $\mu\text{m}$ / $12^\circ$ | $0.37 \pm 0.09^*$ | $121 \pm 33$ | $0.87 \pm 0.25^*$ |
| Natural | $0.14 \pm 0.01$ | $173 \pm 34$ | $0.32 \pm 0.09$ |

### Face recognition

For all pixel sizes and contrast levels, subjects could achieve above 75% accuracy on average, significantly higher than random choice (25%). While faces were nearly instantaneously recognizable with natural vision, more than 5 seconds was needed with simulated vision, since scanning was required to observe all faces due to limited visual field. Increasing number of grey levels and reducing pixel size both improved accuracy and time taken for face recognition. A decrease in pixel size from 100 to 60 μm shortened the response time by around 20% (*p*<0.025 for all contrast levels, 2-sample t-test). There was no significant difference in accuracy between 60 and 100 μm pixels.

## Discussion

Letter acuity and reading speed are the most common metrics for assessment of the quality of vision, especially for low vision patients (31). We added a face recognition task since it is of high priority for patients with atrophic AMD (32). Many psychophysics studies with simulated prosthetic vision were designed to investigate potential capabilities of implants with various numbers of pixels (28, 33–35). Recent clinical results with photovoltaic subretinal prosthesis having 100μm pixels (PRIMA, Pixium Vision) confirmed that prosthetic acuity in AMD patients nearly matches the pixel pitch (19). Moreover, recent measurements with 55 μm pixels in rats demonstrated that grating acuity matches the pixel pitch of this size as well(17). Development of 3-dimensional electrodes enable even smaller pixels, which might provide higher resolution in the future(24). To assess the minimum requirements of a system for restoration of central vision in AMD patients sufficient for reading and face recognition, we decided to evaluate its simulated performance as a function of three parameters: pixel size, field of view (FOV), and number of grey levels.

Previous studies with simulated vision used “phosphenated” images (28, 36, 37). A dot with either a 2D-Gaussian or flat profile was displayed to simulate an activated pixel, while adjacent dots were spaced according to the pixel pitch of the implant, resulting in dark gaps between the simulated phosphenes (38). However, in the PRIMA clinical study, when viewing various line patterns, patients reported perceiving continuous lines, instead of a row of disconnected phosphenes. Therefore, in our study, we used tightly packed pixels, akin to those of a typical consumer monitor, with no dark gaps in between.

Another difference between the current study and previous ones is the choice of FOV. Since other implants were designed for inherited retinal degenerations which cause complete blindness, their functional FOV could be as large as 22°(39). However, geographic atrophy rarely exceeds 4 mm in diameter, and in order to avoid any damage to the adjacent healthy retina, the implant can cover only a part of the scotoma. Hence, subretinal implants for AMD are unlikely to exceed 3 mm in width, corresponding to approximately 10° of the visual angle. In the first feasibility study, the size of the PRIMA implant is 2 mm, corresponding to about 7° of the visual field (19). Therefore, we studied the effect of the FOV on reading speed in the range of 7° to 12°, while all the visual information for a letter acuity or face recognition tasks was packed within 5°of the visual angle.

When our subjects initially were unable to identify the orientation of small Landolt C, they were asked to guess without the experimenter affirming the answer. Typically, the subjects could correctly detect an extra line or two of the acuity chart, which explains their performance exceeding the sampling limit by about 0.2 LogMAR, as can be seen in Figure 3. This strategy is based on scanning the object and identifying a darker or a flickering size of the unresolved blob, which is sufficient for determining the Landolt C orientation. Such strategy can be used for other tasks within a small pool of target patterns, such as letter recognition, but is unlikely to help in identification of unknown objects and patterns.

It was repeatedly shown in the past that accuracy of the face recognition is highly dependent on image resolution, as summarized in (35). With 16×16 phosphenes per face over 9.4° visual field, and 10 levels of grey without scanning, subjects could differentiate faces with up to 84% accuracy (40), one of the highest reported. In another study with 24×24 phosphenes within 18° FOV, accuracy was 65%, and it reached 88% with 32×32 arrays (41). In the current study, focused on modeling small implants in the central macula, we used substantially smaller images (face spanning 5° x 5°) with higher pixel density, while the numbers of pixels per image were comparable to those in previous studies. We found that nearly perfect accuracy can be achieved at 8 grey levels with 60 μm pixels, corresponding to a 24 x 24 grid. On top of using tightly packed pixels and allowing for head scanning, another likely explanation of improved performance is that when the most prominent facial features lie within the fovea (<2 mm in diameter), subjects can spend less effort on scanning, and focus more on evaluating the facial details.

Interestingly, forced-choice face differentiation in our study required significantly fewer pixels than object recognition in a previous study (42). With 100 μm pixels, corresponding to approximately 200 pixels per face, our subjects could differentiate faces at >75% accuracy. This is much less than about 560 pixels needed to recognize objects covering about 10° visual field on a de-cluttered background. The difference could be due to great simplification of the task when a reference is immediately available, compared to naming an object from a large pool of options. Another possibility is that faces could be a surprisingly easy class of images to discern. In a study involving different classes of objects and animals (43), subjects demonstrated >80% recognition rate on all images using 24 x 24 pixels. However, with 16 x 16 pixels, no one could recognize a car, but 90% could identify a dog, which coincides with the accuracy and parameters in our face recognition task. It is also important to keep in mind that in our study the faces we presented on a white background, while with a more cluttered natural background, 2-3 times more pixels maybe needed to achieve the same accuracy (42).

In conclusion, with simulated prosthetic vision in AR glasses, subjects demonstrated letter acuity slightly exceeding the sampling limit, and high efficacy in face recognition even with 100 μm pixels. These results indicate that photovoltaic subretinal implants with 100μm pixels currently available for clinical testing may be helpful for reading and face recognition in patients who lost central vision due to retinal degeneration. As expected, smaller pixels significantly improve visual performance, and therefore, further reduction in pixel size may greatly enhance the outcomes in the future.

## Acknowledgements

Supported by the National Institutes of Health (Grants R01-EY-018608, R01-EY-027786), the Department of Defense (Grant W81XWH-15-1-0009), Stanford Institute of Neuroscience, and Research to Prevent Blindness.

Stimulus images courtesy of Michael J. Tarr, Center for the Neural Basis of Cognition and Department of Psychology, Carnegie Mellon University, http://www.tarrlab.org/. Funding provided by NSF award 0339122.

D.P. is consulting for Pixium Vision. D.P.’s patents related to retinal prostheses are owned by Stanford University and licensed to Pixium Vision. E.H. and J.B. declare no competing financial interests.

## Supplemental Materials

**Supp. Figure 1.** Single-character visual acuity (n=13 for 30 and 60 μm pixels; n=19 for natural vision and 100 μm pixels). In addition to the data presented in Figure 3, the results of ETDRS and Landolt C VA are shown separately here. Error bars are presented in terms of s.d.

