## Supplementary figures and images for "Performance of Complex Visual Tasks using Simulated Prosthetic Vision via Augmented-Reality Glasses"

### Supplemental Figure 1

Supp. Figure 1

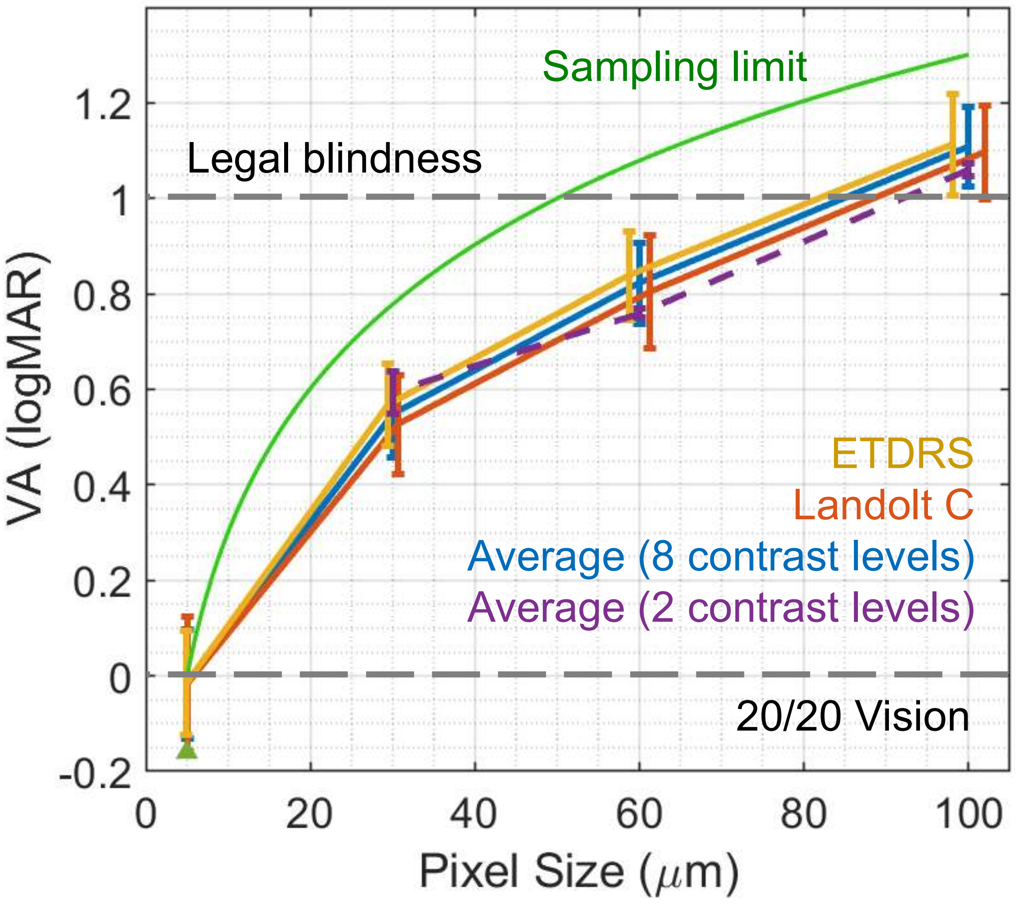
